# An overlooked distal regulatory signal on a shared haplotype drives the neuroinflammatory cascade underlying *GBA1*-associated Parkinson’s disease

**DOI:** 10.64898/2026.07.10.737850

**Authors:** Rahel Feleke, Winston Lau, Philip Zahariev, Bethany Quinton, Alexander Hull, John Hardy, Kerri J. Kinghorn, Dallas M. Swallow, Toby Andrew, Nikolas Maniatis

## Abstract

**BACKGROUND:** The persistent puzzle of how *GBA1* coding variants underlying monogenic Gaucher disease confer such strong risk for polygenic Parkinson’s disease (PD) remains unresolved. Here, we sought to resolve this long-standing paradox.

**METHODS:** Using the Parkinson’s Progression Markers Initiative resource, we integrated whole-genome sequencing from sporadic PD cases and controls with matched transcriptomic, epigenomic and cerebrospinal fluid proteomic data across Northern European and Ashkenazi Jewish populations. Ancestry-specific genetic maps enabled precise integration of disease localisation with eQTL, pQTL and chromatin architecture, linking genetic association to molecular mechanism, while evolutionary genetic analyses resolved the regional haplotypic architecture.

**RESULTS:** We identified a PD-associated regulatory signal located 150 kilobases distal to *GBA1* that robustly replicated across all molecular datasets and both ancestries. Integrative mapping identified rs77268551-G as the candidate regulatory variant, which we show resides on an extended haplotype also carrying the Gaucher disease-causing N370S allele. Evolutionary analyses reveal a clear signature of recent positive selection acting on the Gaucher disease-causing N370S allele, giving rise to this extended shared haplotype that likely concealed the underlying distal PD-associated regulatory signal. rs77268551-G resides within a neuronal enhancer with features consistent with super-enhancer activity that orchestrates hierarchical enhancer-to-enhancer-to-promoter interactions with eleven *cis*-target genes, including *ADAR*, *IL6R* and *GBA1*, driving a coordinated transcriptional programme. Proteomic profiling identified genotype-specific cerebrospinal fluid protein signatures, including PARK7, BIN1 and SAA1, consistent with a neuroinflammatory programme.

**CONCLUSIONS:** The findings support a dual-hit model in which a distal regulatory element is the primary driver of PD-associated molecular programmes, while co-inherited *GBA1* protein-coding variation on the same haplotype modifies and amplifies the risk established by the regulatory variant, giving rise to the clinically distinct *GBA1*-associated PD phenotype. Regulatory activation and *GBA1*-related functional effects converge on neuroinflammatory and lysosomal pathways, and the resulting molecular signatures suggest genotype-linked biomarkers with potential clinical relevance.

## INTRODUCTION

Over the past two decades, mounting evidence has linked Parkinson’s disease (PD) to genetic variation at *GBA1*, which encodes the lysosomal enzyme glucocerebrosidase (GCase). Pathogenic coding variants in *GBA1* cause Gaucher disease, a recessive lysosomal storage disorder characterised by reduced GCase activity and the toxic accumulation of glucocerebrosides within macrophages and neurons. Clinical observations that a subset of individuals with Gaucher disease develop parkinsonism, and that their relatives are at increased risk of PD, first suggested a connection between *GBA1* and PD susceptibility^1^. Since then, numerous *GBA1* variants have been associated with PD^2,3^, including the common N370S (p.N409S) variant, whose frequency varies across populations^4^. This variant is particularly common in Ashkenazi Jewish populations, accounting for ∼15% of PD compared with 3% in non-AJ groups^2,4^. Across ancestries, *GBA1* variants are consistently associated with high PD risk (odds ratio ∼3-8)^2,3^.

Despite the strength and reproducibility of these associations, the mechanism linking *GBA1* variation to sporadic PD remains unclear. Reduced GCase activity has been associated with accumulation of α-synuclein, a key neuropathological feature of PD^5^, but multiple and sometimes conflicting^6^ models have been proposed to explain this relationship^7–9^. Conflicting evidence also exists regarding which pathogenic *GBA1* variants contribute to the clinically distinct *GBA1*-PD phenotype, characterised by earlier onset and more rapid cognitive and motor decline. While more severe variants such as L444P have been linked to worse PD outcomes, other studies report similar progression across *GBA1* carriers regardless of variant type^10,11^. A recent large-scale meta-analysis by the International Parkinson’s Disease Genomics Consortium (IPDGC)^12^ reaffirmed strong association signals at the *GBA1* locus, reinforcing its central importance in PD genetics. However, whether the genotype-phenotype association reflects direct effects of coding variation, broader regulatory mechanisms, or a combination of both, remains unclear.

We investigated the hypothesis that regulatory variation, rather than coding variation alone, contributes to the role of this locus in PD susceptibility. This shifts the focus to identifying genes under regulatory control within this complex genomic region. Such a model is more consistent with the polygenic architecture of sporadic PD and may help explain why decades of focus on *GBA1* coding variation have not yielded a unifying mechanism.

Dissecting this region, however, presents substantial analytical challenges. *GBA1* spans approximately 8 kb on chromosome 1q22 and lies within a gene-dense genomic interval that includes the highly homologous pseudogene *GBAP1* (∼96% sequence identity), as well as *MTX1* and its pseudogene *MTX1P*. This architecture promotes gene conversion events^13,14^ and local genomic rearrangements^15^, while founder effects and demographic history since the Jewish Diaspora have generated extended linkage disequilibrium (LD) across the region^16^. The interplay of these features create complex and population-specific LD patterns that complicate fine mapping and hinder the distinction between coding and regulatory effects.

To resolve this regional complexity, we built LD maps that capture the combined influences of recombination, demography and gene conversion. Using the multi-layered PPMI resource (Parkinson’s Progression Markers Initiative, https://www.ppmi-info.org), we integrated these population-specific maps with PD-specific multi-omics data across Ashkenazi Jewish (AJ) and Northern European (nEUR) populations. Through this integrative approach, we identify a previously unrecognised distal regulatory signal located 150 kb upstream of *GBA1* that resides on a shared haplotype with one of the most common *GBA1* variants, N370S. These findings resolve the longstanding discrepancy between the strong genetic association of *GBA1* variants with PD and the absence of a coherent mechanistic explanation. By linking this regulatory signal to coordinated transcriptional, chromatin architectural, and proteomic changes, our study provides a mechanistic explanation for how variation at this genomic region contributes to disease risk and the clinically distinct *GBA1*-associated PD (*GBA1*-PD) phenotype, reframing the role of *GBA1* within a broader haplotypic, regulatory and inflammatory framework.

## METHODS

### Study design and overview

We first applied our localisation framework to meta-analysed summary statistics from 16 genome-wide association studies made publicly available by the IPDGC^12^, comprising approximately 54,000 sporadic PD cases and 846,000 controls. The novel PD localisation signal identified in this discovery analysis motivated an in-depth integrative investigation of the region using genomic, transcriptomic, three-dimensional genomic, epigenomic, and proteomic data generated by the PPMI (https://www.ppmi-info.org). PPMI was selected as the principal resource because it uniquely combines multiple molecular layers within a disease-specific cohort, enabling integrative fine-mapping, cross-ancestry replication, and direct comparison of genetic and molecular signals within a unified analytical framework (Figure 1A).

**Figure 1.**
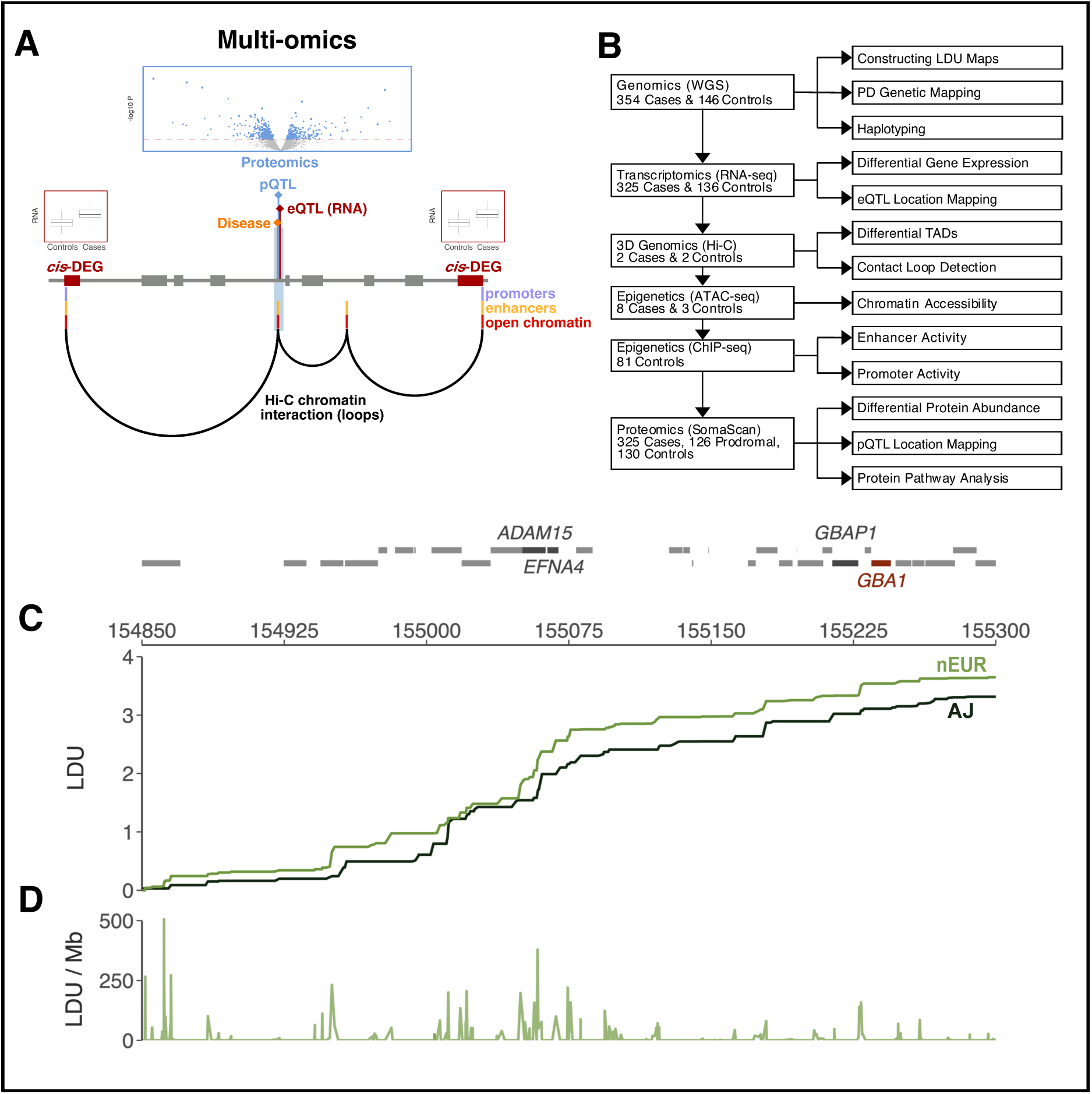
Overview of study design and integrative mapping framework using genetic LDU maps. **(A)** Schematic diagram of the multi-omics PD architecture investigated in this study. Disease mapping in orange, eQTL (expression) mapping in dark red, and pQTL (protein) mapping in blue. All three signals collocate at a region of enhancer activity with differentially open chromatin at this site. This serves as the anchor point for a regulatory network identified using Hi-C interaction data (in black). This network implicates the *cis* genes with collocating eQTL at the PD locus, which are also differentially expressed (DEGs) between PD cases and controls. **(B)** Analytical workflow of the integrative mapping approach, showing the sequential analytical stages across omics data together with their corresponding sample sizes. **(C)** Cumulative linkage disequilibrium unit (LDU) genetic maps in nEUR and AJ population constructed from the PPMI data, with extended LD blocks observed as plateaus, separated by stepwise increases corresponding to LD breakdowns (observed as spikes in (D)). **(D)** LD breakdowns across the *GBA1* region identified using nEUR LDU maps, plotted as LDU per megabase (LDU/Mb).

### Study populations and data

PPMI is a multicentre longitudinal and deeply phenotyped study designed to evaluate clinical, genetic, biological, and molecular data across all stages of PD. Full study protocols, standard operating procedures, and ethics documentation are available at https://www.ppmi-info.org/study-design/archive-of-research-docs-and-sops.html. We used data from the Clinical cohort, which includes participants with whole-genome sequencing (WGS), whole-blood RNA sequencing (RNA-seq), induced pluripotent stem cell (iPSC)-derived dopaminergic neurons for Hi-C and ATAC-seq assays, and cerebrospinal fluid (CSF) for proteomics (Figure 1A).

Population structure was assessed using multidimensional scaling against the 1000 Genomes reference panel and validated using ADMIXTURE (Supplementary Figures S1–S2). Two genetically distinct clusters corresponding to nEUR and AJ ancestry were identified and analysed separately. The AJ cluster was corroborated by near-perfect overlap with self-reported AJ ancestry, consistent with prior evidence that individuals of AJ ancestry form a genetically distinct cluster even when derived from a single Jewish grandparent^17^.

Participants were classified as PD cases, prodromal individuals, or healthy controls based on clinical assessment, imaging, and genetic screening. Cases and controls were matched for age and sex, and prodromal individuals were excluded from all analyses except proteomics (Supplementary Figure S3). *GBA1* carrier status (*GBA1*^+^) was defined based on the presence of pathogenic variants and used to identify individuals with the *GBA1*-PD clinical phenotype. Carrier status in PPMI was determined by Sanger sequencing of known pathogenic variants (including N370S, R502C, R159W, L29Afs*18, IVS2+1G>A, L444P, and R535H), an essential step given the high sequence homology between *GBA1* and its pseudogene *GBAP1*^13,14^.

Among *GBA1*^+^ carriers, N370S was overwhelmingly predominant, accounting for 92% of AJ and 63% of nEUR carriers (the latter based on 8 individuals), consistent with its established role as the most frequent variant associated with *GBA1*-PD phenotype^2^. Figure 1B summarises the number of participants by ancestry, analysis, and omics platform. As participants did not contribute to every assay (for example, Hi-C data were available for a subset only), sample sizes varied across analyses. A detailed breakdown of sample sizes for each analysis is provided in Supplementary Table 1.

### Analytical framework and genetic localisation

The mapping framework was designed to require convergence of multiple independent lines of evidence on a shared, narrowly localised interval. Co-localisation was defined stringently as the convergence of signals from independent omics platforms within a few kilobases of a shared inferred causal location, rather than the much broader intervals commonly used in the field, and was required to replicate independently across both AJ and nEUR populations. To achieve this resolution, mapping was performed using population-specific genetic maps, in which each SNP is assigned a genetic location expressed in linkage disequilibrium units (LDUs)^18,19^. LDU maps capture regional haplotype structure by modelling the cumulative breakdown of LD across the locus. Figure 1C shows LDU maps constructed separately for nEUR and AJ PPMI participants, revealing plateaus of conserved LD separated by regions of LD breakdown, including the discontinuity observed between *GBA1* and *GBAP1*. These patterns reflect not only recombination but also other evolutionary forces, including demographic history and gene conversion, processes known to shape the complex architecture of the *GBA1* region^13–16^.

Association testing was conducted using a multi-locus composite-likelihood framework that jointly models information across neighbouring SNPs within the 450 kb analytical window together with their LDU distances. Because each SNP is assigned a population-specific LDU position, local LD structure is incorporated directly into the localisation model rather than considered retrospectively. The approach estimates the most likely causal location and its support interval within the analytical window localisation^18,19^. The resulting likelihood-deviance profiles are visualised as localisation curves, enabling direct comparison of disease, transcriptomic, epigenomic, chromatin, and proteomic signals within a common genetic coordinate system. Although all localisation analyses were performed in LDU space, localisation curves are displayed against physical genomic coordinates (kb) to facilitate visual comparison across molecular datasets and with established genomic annotations. For the initial IPDGC analysis, summary statistics for all published markers^12^ included imputed SNPs were used before performing multi-locus localisation. However, all subsequent analyses relied exclusively on directly observed genotypes, thereby avoiding the potential limitations of genotype imputation for disease localisation, as previously demonstrated^20^. Further details are provided in the Supplementary Methods.

### Haplotype and evolutionary analyses

To determine whether the distal association signal segregated independently of *GBA1*, phased haplotypes spanning N370S and the lead variant were reconstructed and visualised using neighbour-joining clustering. Extended haplotype homozygosity (EHH) analysis was used to assess evidence for recent positive selection. Because PPMI is clinically ascertained, EHH findings were replicated in the population-based UK Biobank to distinguish selective effects from demographic enrichment (Supplementary Figure S5). Further methodological details are provided in the Supplementary methods.

### Transcriptomic analyses

Bulk RNA sequencing data from PPMI whole blood were analysed in participants with paired WGS and transcriptomic data. Expression data were obtained for all 174 genes within a 3 Mb *cis*-region surrounding *GBA1* (±1.5 Mb). Two complementary analyses were performed. Differential expression (DEG analysis) between PD cases and controls was assessed using negative binomial regression, adjusting for demographic and technical covariates. Expression quantitative trait locus (eQTL) mapping tested associations between SNPs within the same 450 kb analytical window used for disease mapping and gene expression using the same modelling framework as disease mapping, thereby providing a regulatory location estimate and corresponding significance test for each gene.

Genes were considered PD-relevant *cis-*targets only if they satisfied stringent criteria: (i) eQTL localisation overlapped precisely with the PD-associated interval; (ii) the gene was differentially expressed (FDR <0.05); and (iii) results replicated independently in both AJ and nEUR populations. External validation was performed using independent neuronal transcriptomic datasets^21–26^. Full details are provided in the Supplementary methods.

### Three-dimensional genomic and epigenetic analyses

Chromatin architecture was investigated using Hi-C data from PPMI iPSC-derived dopaminergic neurons, generated by the FOUNDIN-PD project^27^. Hi-C contact matrices were used to identify topologically associating domains (TADs) and chromatin interaction loops across the 3 Mb *cis-*region.

Loop detection was performed at high resolution to identify enhancer–promoter interactions, with a focus on contacts originating from the disease-associated locus. Chromatin accessibility was assessed using single-nucleus ATAC-seq data from the same cellular system. Differential accessibility analyses were performed comparing carriers and non-carriers of the lead variant. Chromatin state annotation was obtained from publicly available neuronal epigenomic datasets and used to identify enhancer elements within the region of interest. Detailed processing steps are provided in the Supplementary Methods.

### Proteomic analyses

CSF proteomic data were obtained from the curated PPMI Proteomics Working Group dataset for drug-naïve PD and prodromal participants with matched genotype data. Protein abundance was quantified using the SomaScan platform. Generalised linear models were used, adjusting for demographic variables and principal components derived from the proteomic matrix. Differentially abundant proteins were defined after FDR correction. Inclusion of prodromal participants enabled assessment of potential early disease-associated proteomic changes.

For proteins showing evidence of differential abundance, protein quantitative trait locus (pQTL) mapping was performed using the same localisation framework. These analyses assessed whether *trans*-proteomic signals co-localised with both the PD and *cis*-eQTL intervals, thereby determining whether genetic effects at this locus extend from transcriptional regulation to the proteome. Given the modest sample size of the proteomic dataset, which was further reduced by stratification, analyses were performed separately in nEUR and AJ populations. Only proteins showing nominal evidence of differential abundance in both populations were carried forward to combined meta-analysis. Genome-wide pathway enrichment analyses of differentially abundant proteins were performed using KEGG (adjusted P<0.05). Additional analytical details are provided in the Supplementary Methods.

## RESULTS

### Fine mapping localises PD risk to a regulatory interval distal to *GBA1*

We first applied our localisation framework to meta-analysed summary statistics from 16 genome-wide association studies made directly available by the IPDGC^12^, comprising 53,858 sporadic PD cases and 846,380 controls of European ancestry (Figure 2A, B). Contrary to expectations, the signal of causality did not map to *GBA1* itself but instead mapped approximately 150 kb upstream, within a sharply delineated 30 kb support interval at a location proximal to *EFNA4*. The localisation curve shows the estimated causal location as the deepest point, representing the best fit for the multi-locus model, with a narrow support interval reflecting high mapping precision and clear separation from *GBA1* (Figure 2B).

**Figure 2.**
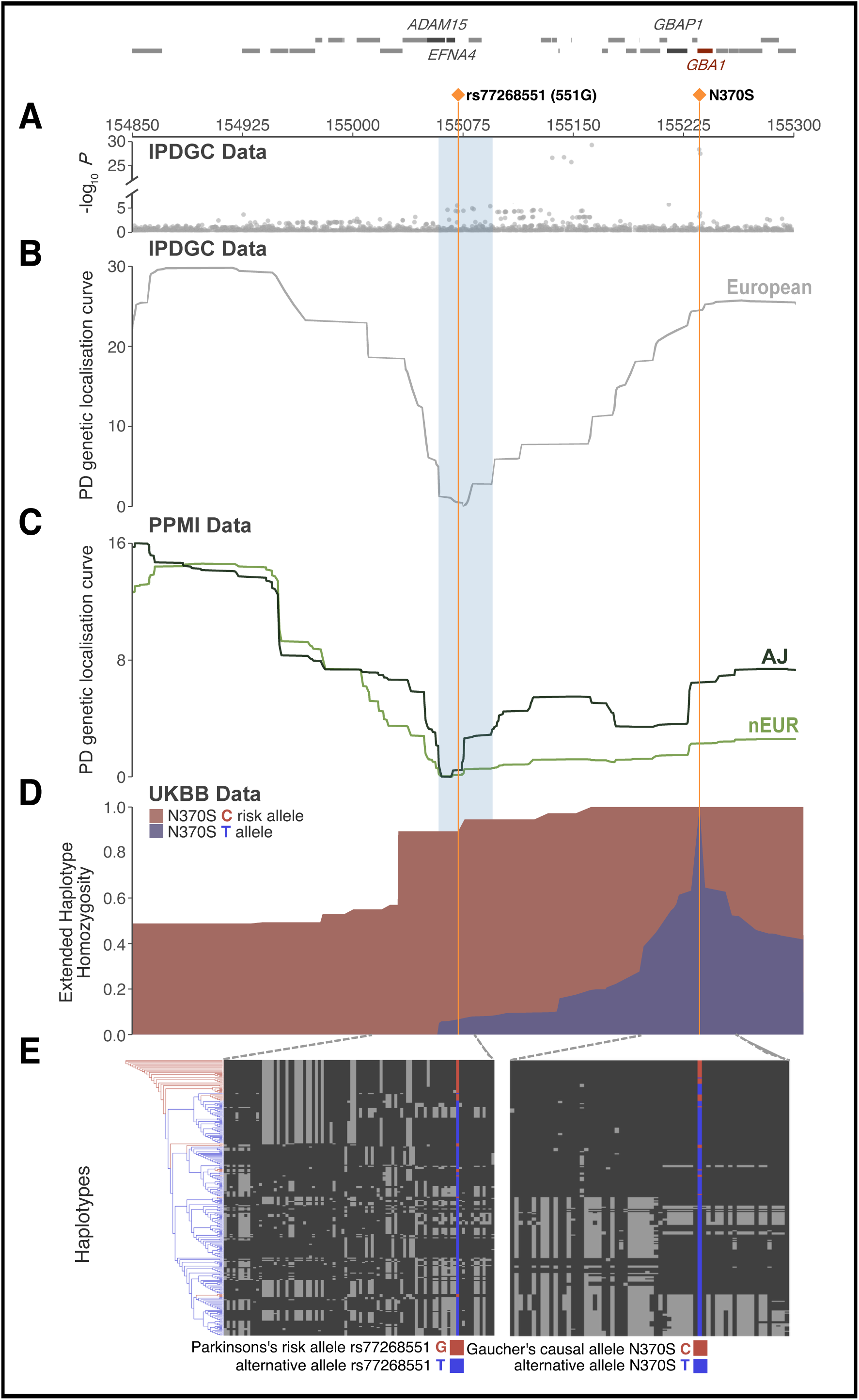
Discovery and validation of the distal PD regulatory interval and its shared haplotype with *GBA1*. **(A)** Single-SNP association results from the IPDGC GWAS meta-analysis across the region. **(B)** Initial localisation curve from IPDGC meta-data using a multi-locus mapping framework based on the nEUR genetic LDU map. The Y-axis shows the deviance from the best-supported causal location, with lower values indicating stronger statistical support. The left orange line marks the deepest point of the localisation curve, representing the inferred causal location for PD (*P*=5.9E-22), the surrounding blue shaded region indicates the 30 kb support interval. The right orange line marks the location of the *GBA1* N370S variant. **(C)** Independent localisation curves using the PPMI data for nEUR and AJ populations for *GBA1*^+^ PD cases vs controls, both converging precisely on the same narrow interval encompassing the candidate functional variant rs77268551 (551G); blue shading indicates support interval. **(D)** Extended Haplotype Homozygosity (EHH) analysis of the *GBA1* N370S variant in AJ individuals from the population-based UK Biobank, showing slower decay of homozygosity for the Gaucher’s-associated risk allele (C) compared with the alternative allele. The extended haplotype carrying the risk allele spans the 551G region, consistent with a recent selective sweep and a long-range haplotype conservation. **(E)** Phased Haplotype structure demonstrating that the risk allele of *GBA1* N370S (red, right) travels on the same extended haplotype as the PD risk allele G of 551G (red, left), whereas the alternative alleles at both loci are shown in blue. Black background denotes extended haplotypes, grey denotes breakdown in LD.

Inspection of the published IPDGC summary statistics reveals that this *EFNA4* interval harbours multiple SNPs nominally associated with PD (Figure 2A), despite being flanked by recombination hotspots (see *EFNA4* in Figure 1D). In contrast, *GBA1* contains far fewer even if more significant, SNPs despite lying within a region of extended LD (Figure 1C, D), where a greater number of associated SNPs would typically be expected^20^.

To replicate this unexpectedly distinct localisation estimate, we next analysed PD cases and controls of nEUR and AJ ancestry in the PPMI cohort separately, applying the same LDU-based mapping strategy using genetic maps constructed for each ancestry. This enabled validation across populations while accounting for ancestry-specific differences in LD architecture (Figure 2C). Initial analyses of all cases and controls yielded no significant results. Instead, precise and significant localisation was achieved only when analyses were restricted to PD cases with the *GBA1*-PD phenotype (i.e. patients carrying a functional *GBA1* risk allele, accurately defined by Sanger sequencing). Among *GBA1^+^* carriers, N370S was the predominant Gaucher disease-causing variant. Stratifying by *GBA1^+^* status and comparing these patients with all controls yielded strikingly concordant localisation profiles for both nEUR and AJ populations (*P*=9.8E-12 and 1.7E-18, respectively). Importantly, both signals co-localised independently within 4 kb of the IPDGC discovery location. The larger deviance values (Y-axis) observed in IPDGC localisation curve relative to PPMI reflect the substantially greater sample size of IPDGC, which increases sensitivity to deviations from the best-fit causal location.

Within this narrow 30 kb support interval, we identified three candidate non-coding variants located approximately 10 kb apart, with rs77268551 positioned centrally between the other two (Table 1). The risk alleles of these regulatory variants were markedly more common in *GBA1*^+^ PD cases than in controls, with frequencies of 50% versus 6% in AJ and 31% versus 2% in nEUR for rs77268551, respectively (Table 1). These differences translated into exceptionally high odds ratios (21.0 in nEUR and 15.0 in AJ). We therefore focused on rs77268551 as the lead candidate variant, hereafter referred to as 551G, because it consistently aligned with the best-fit location estimate across all three localisation curves (PPMI EUR, PPMI AJ, and IPDGC EUR). Subsequent stratification analyses were therefore based on carrier status of the lead variant rs77268551-G rather than *GBA1*^+^ status.

**Table 1.** Allele frequencies and association of variants at the distal PD regulatory signal and *GBA1*. A summary of the lead variants, including their risk allele frequencies (RAF) and association statistics in European (nEUR) and Ashkenazi Jewish (AJ) populations. For each population, analyses are presented for all PD cases versus controls and for analyses restricted to *GBA1* positive (*GBA1*^+^) PD cases versus controls. Odds ratios and *P*-values are reported for each variant.

| Unstratified cases vs controls |  |  |  |  | Ashkenazi Jewish (AJ) |  |  |  | Northern Europeans (nEUR) |  |  |  |
| --- | --- | --- | --- | --- | --- | --- | --- | --- | --- | --- | --- | --- |
| Closest gene | SNP | Position Bp | Allele Risk | alt | Frequency % of risk allele |  | OR | P | Frequency % of risk allele |  | OR | P |
|  |  |  |  |  | Case | Ctrl |  |  | Case | Ctrl |  |  |
| *ADAM15* | rs11589479 | 155060832 | A | G | 34.7 | 20.6 | 2.0 | 0.10 | 15.9 | 16.5 | 1.0 | 0.82 |
| *EFNA4* | rs77268551 (551G) | 155071721 | G | T | 20.2 | 6.2 | 3.8 | 0.06 | 3.0 | 2.0 | 1.5 | 0.46 |
| *EFNA3* | rs140309215 | 155082256 | C | G | 17.7 | 3.1 | 6.7 | 3.5E-02 | 2.5 | 2.0 | 1.3 | 0.66 |
| *GBA1* | rs76763715 (N370S) | 155235843 | C | T | 15.5 | 0.0 | 6.6 | 0.09 | 1.1 | 0.0 | 3.4 | 0.67 |
| Stratified cases (*GBA1*<sup>+</sup> only) vs controls |  |  |  |  |  |  |  |  |  |  |  |  |
| *ADAM15* | rs11589479 | 155060832 | A | G | 62.9 | 20.6 | 6.5 | 5.2E-05 | 50.0 | 16.5 | 5.0 | 3.2E-03 |
| *EFNA4* | rs77268551 (551G) | 155071721 | G | T | 50.0 | 6.2 | 15.0 | 7.1E-05 | 31.2 | 2.0 | 21.0 | 9.6E-05 |
| *EFNA3* | rs140309215 | 155082256 | C | G | 48.5 | 3.1 | 29.2 | 7.8E-06 | 28.6 | 2.0 | 18.8 | 5.6E-04 |

### Haplotype analysis reveals co-segregation of 551G with the *GBA1* N370S allele

Complete linkage disequilibrium (D′ = 1) between the 551G and N370S variants was observed in the unstratified nEUR and AJ datasets. In parallel, stratification by *GBA1*^+^ status revealed a marked increase in the effect size (OR) of the 551G signal. These observations point to shared inheritance on a common haplotype. To test this directly, phased haplotypes spanning N370S and 551G were reconstructed and visualised using neighbour-joining clustering (Figure 2D and 2E). Extended haplotypes were observed only in individuals carrying both the G (551G) and C (N370S) risk alleles (Figure 2E, shown in red), whereas individuals carrying the alternative alleles showed substantially greater haplotypic diversity and breakdown of extended haplotype structure across the surrounding region (Figure 2E, shown in blue). This finding was further supported by adapting the population differentiation metric F_st_. Rather than comparing populations, we quantified genetic differentiation between *GBA1*^+^ and non-carriers for each SNP within each ancestry. Elevated F_st_ across the same extended haplotypic interval in both populations (Supplementary material) also demonstrated marked differentiation associated with the shared haplotype.

These haplotypes clearly show that the Gaucher disease-causing N370S allele C co-segregated almost completely with the G allele of 551G, indicating that the distal regulatory variant has travelled alongside N370S on a shared haplotypic background. This shared haplotypic background may have obscured the underlying regulatory signal at 551G, complicating efforts to distinguish functional effects related to Gaucher disease from those driven by distal regulatory variation in PD.

### Extended haplotype analysis indicates recent positive selection at the *GBA1* N370S allele

What gave rise to such an extended haplotypic structure? We next examined extended haplotype homozygosity (EHH) as evidence of recent positive selection, focusing on 1,388 individuals of AJ ancestry from the UK Biobank to avoid disease-related ascertainment bias (Figure 2D). Using the *GBA1* N370S variant as the focal marker, EHH analysis revealed that haplotypes carrying the Gaucher disease-causing C allele (shown in red) decayed markedly more slowly than those carrying the alternative T allele (shown in blue), a pattern consistent with a recent selective sweep (Figure 2D).

Accordingly, carriers of the N370S C allele share unusually long haplotypes compared with non-carriers, reflecting reduced historical recombination across the region encompassing 551G. To confirm that this signal was not specific to the UK Biobank cohort, we replicated the analysis in the clinically characterised PPMI dataset, where the same pattern was observed (Supplementary Figure S5). These findings support the conclusion that this genomic region bears a clear signature of recent positive selection, suggesting that a strong historical selective force has shaped and preserved this haplotypic architecture.

### Co-localisation of PD and *cis*-eQTL signals reveals an inflammatory regulatory network at 551G

Having established that PD risk maps distal to *GBA1*, we next investigated whether genetic variation within the 551G interval exerts regulatory effects on gene expression and, in doing so, re-examined the role of *GBA1* itself. A central question was whether the longstanding association between *GBA1* and PD arises solely from the extended haplotypic background shaped by the selective sweep, or whether *GBA1* is also functionally implicated as a target *cis-*gene under direct regulatory control of the 551G locus.

We therefore examined whether variation within the 551G interval acts as a *cis*-regulatory interval (eQTL) influencing expression of nearby genes and, if so, whether those same genes are differentially expressed in PD. To address this, we analysed individuals with complete genomic and whole-blood RNA-sequencing data from PPMI. Genes were considered PD-relevant *cis*-targets only if they satisfied the predefined criteria described in the Methods, requiring co-localisation of both PD and eQTL signals together with concordant differential expression.

Analyses of transcriptomic data across the 174 genes in the 3-Mb *cis-*region flanking *GBA1* identified eleven *cis-*genes satisfying all criteria. Localisation curves constructed independently for AJ and nEUR populations, and meta-analysed across the eleven *cis-*genes (Figures S6 and S7, respectively), showed convergence of regulatory locations at the same 551G interval that defines the PD association (Figure 3A), demonstrating replication across ancestries. When analyses were repeated using all PD cases irrespective of 551G carrier status, no significant eQTL localisation was detected, indicating that the regulatory signal is specific to the *GBA1*-PD subtype and becomes diluted by genetic heterogeneity when all samples are combined.

**Figure 3.**
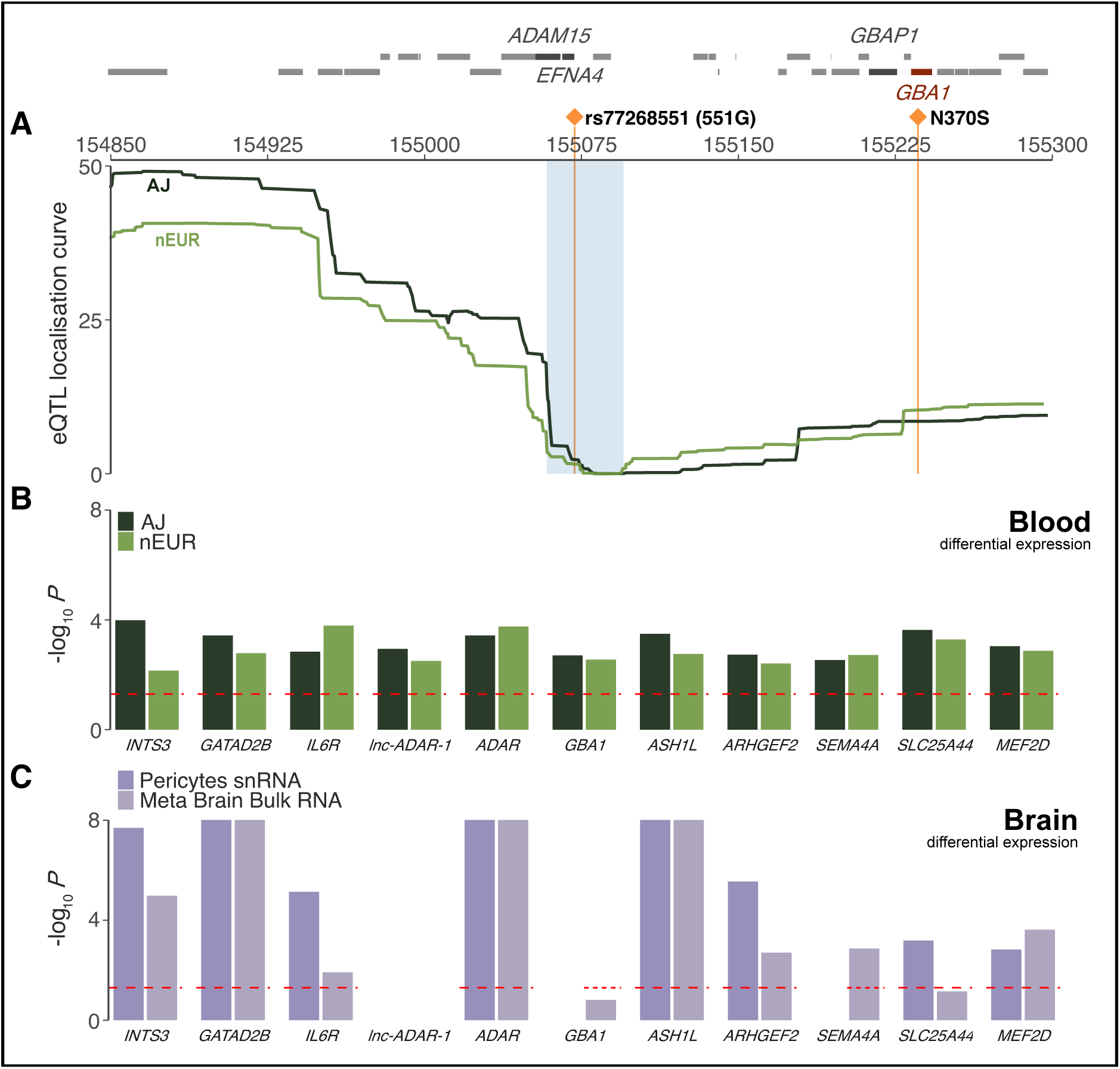
Fine scale eQTL mapping links 551G to coordinated *cis-r*egulation across blood and neuronal cells. **(A)** Meta-analysed blood eQTL localisation curves across eleven *cis-*genes converge within the narrow 551G interval (blue shading) in both AJ (dark green) and nEUR (green) populations. Individual *cis-*gene localisation curves are shown in the Supplementary Figures S6 and S7. **(B)** Differential expression analyses (all cases vs controls) for whole-blood shows all eleven *cis-*genes as differentially expressed (red dashed line denotes nominal significance). All genes are upregulated in PD cases compared with controls. **(C)** Replication of the eleven *cis-*DEGs in independent neuronal datasets (pericytes single-nucleus RNA-seq and brain bulk RNA-seq meta-analysis). Pericyte data yielded concordant upregulation with PPMI blood, whereas effect directions varied in brain bulk RNA-seq, consistent with cellular heterogeneity. Absence of a dashed red line indicates missing data.

All eleven *cis-*genes were also significantly differentially expressed in blood between PD cases and controls, with results replicated across both populations (Figure 3B). Differentially expressed gene (DEG) analyses using either all PD cases or only 551G carriers identified the same gene set, with larger effect sizes observed in the carrier-restricted analysis (Supplementary Table S2); for simplicity, 3B presents results from all PD cases. All eleven *cis-*genes showed concordant upregulation in PD cases across populations (Figure 3A & 3B). Because these target genes show evidence for both co-localising eQTLs and disease-associated differential expression, they are referred to here as *cis-*DEGs. Notably, *GBA1* itself was among the eleven *cis-*DEGs, indicating that its upregulation originates from variants within the disease location interval rather than from direct coding effects. These findings suggest a revised interpretation of the *GBA1* region, positioning *GBA1* within a broader inflammatory network under distal regulatory control mediated by 551G. The same distal locus also regulates genes central to immune signalling and inflammation, including *ADAR* and its adjacent lncRNA, *IL6R, SLC25A44, MEF2D*, *ARHGEF2, INTS3, SEMA4A, GATAD2B*, and *ASH1L*. All *cis-*DEGs showed concordant upregulation in PD cases across populations (Figure 3A & 3B), supporting the presence of a coordinated inflammatory transcriptional response.

### Cross-tissue replication reinforces neuroinflammatory signalling at 551G

The uniform upregulation of all eleven *cis-*DEGs in blood prompted independent validation using neuronal transcriptomic datasets for PD (Figure 3C). These included published values from single-nucleus RNA-seq of pericytes^21^ and our own meta-analysis of five brain bulk RNA-seq GEO datasets^22–26^. With the exception of three genes with missing data (*lnc-ADAR-1* in both datasets, and *GBA1* and *SEMA4A* in pericytes), all eight remaining genes in pericytes showed replicated upregulation, achieving complete concordance with the PPMI blood signal. In the bulk brain RNA-seq meta-analysis, only two genes (*GBA1* and *SLC25A44*) did not meet the significance threshold (red dashed line, Figure 3C). Pericyte-specific expression mirrored the direction of effect observed in PPMI blood across all evaluable *cis*-genes, whereas bulk brain RNA-seq showed more variable effects, consistent with cellular heterogeneity (Supplementary Table 5).

The lack of differential expression for *GBA1* in bulk RNA-seq meta-analysis likely reflects technical limitations of short-read sequencing^28^. Due to high sequence homology between *GBA1* and *GBAP1*, a substantial proportion of reads may fail to map uniquely and are therefore discarded, leading to underestimation of true *GBA1* expression. This limitation is supported by long-read RNA-seq data^28^ demonstrating higher *GBA1* expression relative to its pseudogene, suggesting that multimapping artefacts may obscure genuine differential expression in short-read datasets.

These results demonstrate that the *EFNA4* region, with 551G as the leading candidate causal variant, drives a coordinated regulatory programme affecting eleven genes across both peripheral and central systems. The concordant upregulation observed in blood and pericyte-specific expression points to a neuroinflammatory programme in which blood brain barrier-associated cell types may play a central role; a concept developed further in the discussion.

### Allele-specific 3D chromatin organisation at 551G reveals enhancer-promoter regulatory interactions

To investigate the regulatory architecture of the 551G locus, we analysed Hi-C chromatin conformation data from iPSC-derived dopaminergic neurons. Data were available from two PD patients, one carrying the 551G risk allele and one carrying the alternative T allele, and from two healthy controls, both of whom carried the T allele (Figure 4). Critically, the G-allele PD patient did not carry the *GBA1* N370S variant, providing a genetically clean separation of regulatory effects attributable specifically to 551G. Across the 3-Mb *cis-*region, we observed allele-specific differences in topologically associating domain (TAD) organisation and chromatin looping between the PD G-allele carrier and the PD T-allele carrier (*P*=0.002; Figure 4A-B), corroborated by concordant changes in chromatin accessibility assessed independently by snATAC-seq.

**Figure 4.**
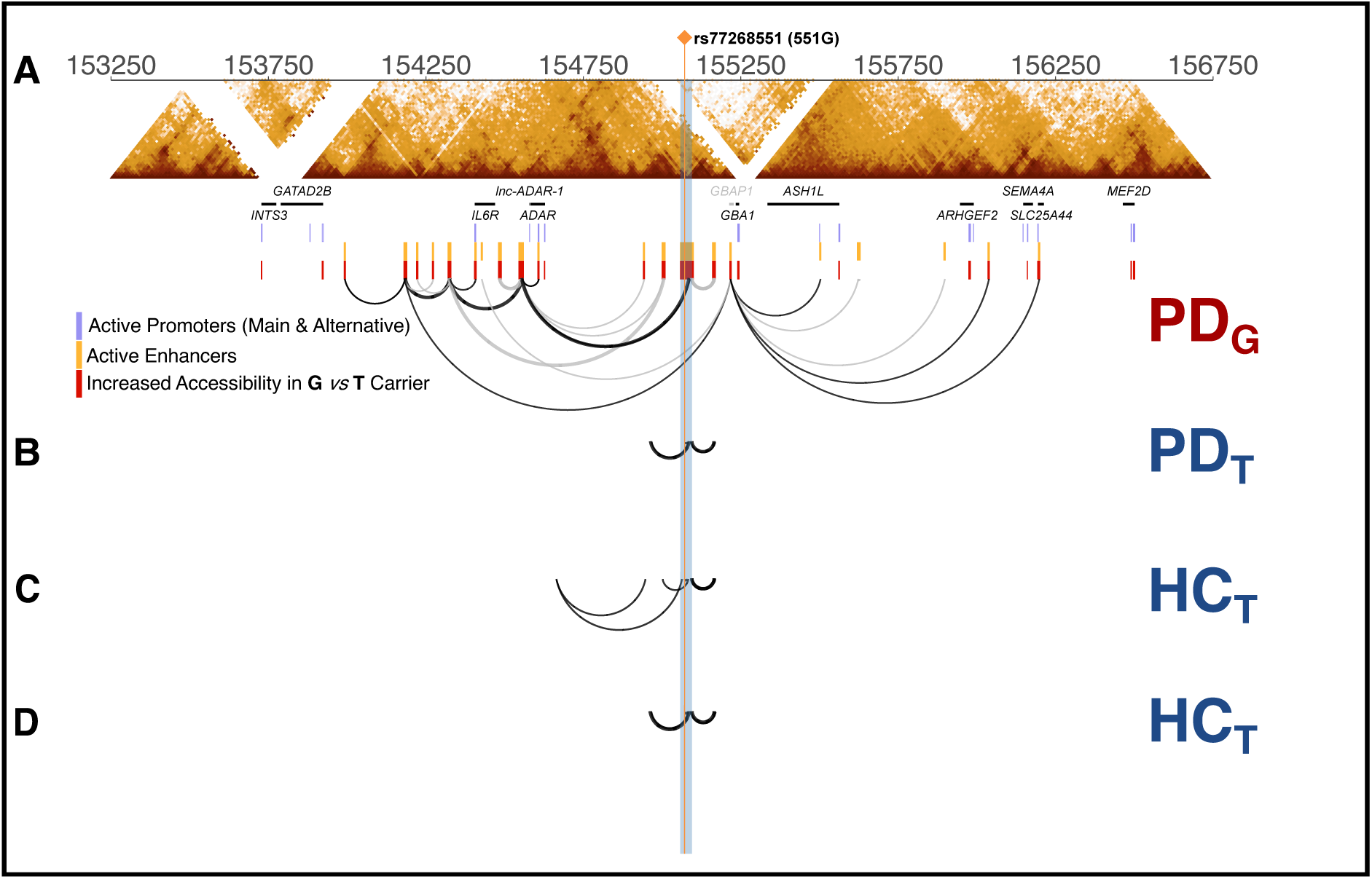
Allele-specific chromatin architecture links the 551G enhancer to *cis-*gene regulation. **(A)** Hi-C data from dopaminergic neurons derived from iPSCs. The upper panel shows topologically associating domains (TADs) across the 3 Mb *cis-*region. The lower panel depicts a hierarchical network of chromatin interactions emanating from the enhancer harbouring 551G (Yellow), which interacts with distal enhancers (yellow) and ultimately anchors at promoter sites (green) of the eleven *cis-*DEGs. These terminal promoter anchors coincide with regions of increased chromatin accessibility (red), as assessed by ATAC-seq, in the 551G (G-allele) carrier compared with the T-allele carrier. This multilayered enhancer–promoter architecture is observed only in the PD patient carrying the G risk allele. **(B)** Hi-C maps from the PD T-allele carrier and **(C, D)** two healthy control T (HC_T_) carriers show simplified chromatin looping and the absence of the hierarchical enhancer-promoter network observed in the 551G carrier.

The narrow interval encompassing 551G is enriched for neuronal enhancer elements, and the 551G variant itself resides within a 1-kb neuronal enhancer^29^, supported by tissue-specific chromatin state annotation (Supplementary Figure S8). Chromatin loops were then traced from the 551G interval to determine whether their terminal anchors mapped to promoters of the 551G-associated *cis*-DEGs.

Indeed, in the PD patient carrying the 551G risk allele, the enhancer formed primary loops with distal enhancers that acted as intermediaries for further enhancer-enhancer interactions before anchoring at promoters of the 551G-associated *cis-*DEGs (Figure 4A). For example, the focal enhancer contacted *ADAR* via a secondary enhancer, whereas *IL6R* required an additional looping step from the same secondary enhancer before promoter contact. Despite the small sample size, this pattern in Figure 4A reflects all chromatin loops emanating from the 551G enhancer interval without any selection based on the *cis*-DEGs identified in this study. The fact that these loops anchor exclusively at these *cis*-DEG promoters, and not at any other genes within the surrounding highly gene-rich region, emerges as an unbiased result that strengthens the regulatory connection.

The observed interaction pattern defines a multi-connected regulatory hub linking 551G to its downstream *cis-*DEG promoters, reflecting an allele-specific mechanism of hierarchical enhancer-promoter communication driven by the PD-associated regulatory signal. By contrast, this architecture was absent in the PD patient carrying the T allele and in healthy controls (Figure 4B-D).

*In silico* motif analysis further indicated that the 551G allele disrupts binding motifs for transcription factors involved in chromatin remodelling and inflammatory signalling, including BPTF, FOXD3, and FUBP1^30–34^ (Supplementary material). When all the findings are viewed together, they clearly point to a nested enhancer network that remodels local 3D genome architecture, which explains the activation of a broader transcriptional programme encompassing the eleven *cis-*DEGs. To assess whether this remodelling is accompanied by changes in chromatin accessibility, we analysed snATAC-seq from iPSC-derived neurons of PD patients. Comparing G-versus T-allele carriers revealed significantly increased accessibility (*P.adj*<0.05) at all enhancer-enhancer interaction sites and at the corresponding promoter anchors, with uniformly greater accessibility in G-allele carriers (highlighted in red in Figure 4A). Although Hi-C coverage was unavailable for a small subset of *cis-*DEGs (*GATAD2B, INTS3*, and *GBA1*), their promoters overlapped precisely with differentially accessible regions. These results provide evidence for allele-specific chromatin remodelling in which the 551G locus drives a functional hierarchical enhancer-promoter network that activates a coordinated inflammatory transcriptional programme.

### Allele-specific proteomic signatures linked to the 551G

We next asked whether the regulatory effects observed at the transcriptomic level extend to the proteome, using CSF data from the same PPMI cohort. Specifically, we examined whether stratifying PD patients by genotype at 551G reveals inflammatory protein signatures consistent with the transcriptomic dysregulation identified above, thereby linking upstream genetic regulation to downstream proteomic changes within the central nervous system.

Protein abundance was compared between 551G carriers (GG or GT) and non-carriers (TT), first in PD cases alone and subsequently after inclusion of prodromal cases to explore early disease signatures and biomarker potential. To prioritise robust signals, analyses focused on proteins showing evidence of association in both populations and exhibiting a pQTL localising to the 551G interval, before combined analyses across replicated proteins.

The CSF proteomic data were obtained from the PPMI Proteomics Working Group for both PD and prodromal participants with matched genotype data. Protein abundance was quantified using the SomaScan platform and compared between 551G carriers (GG or GT) and non-carriers (TT) using generalised linear models. This design isolated genotype-dependent proteomic effects while leveraging the relative proteomic homogeneity of drug-naïve PPMI patients. Protein quantitative trait locus (pQTL) mapping was performed using the same LDU-based localisation framework to assess co-localisation with PD and eQTL intervals. Proteins showing significant differential abundance after FDR correction that also exhibited a pQTL localising to the 551G interval were analysed further. First in PD cases alone and subsequently after inclusion of prodromal cases to explore early disease signatures and biomarker potential. Analyses were conducted separately in nEUR and AJ cohorts, and only proteins nominally significant in both populations were carried forward to a combined analysis.

A total of 22 proteins replicated independently across both ancestries (highlighted in red in Figure 5A and listed in the Supplementary Table 6). Six of these proteins (labelled in Figure 5A) also yielded significant pQTLs that co-localised precisely with the 551G interval (Figure 5B), thus directly linking the same locus that drives PD risk and gene expression to downstream protein abundance. Among these six proteins with colocalising pQTLs, PARK7, SAA1, BMP15, and LIF were elevated in 551G carriers. Elevated PARK7 (DJ-1) is consistent with a response to inflammatory or oxidative stress associated with the 551G regulatory programme, while elevated SAA1 reinforces the transcriptomic evidence for a pro-inflammatory state and links the 551G locus to neuroinflammatory processes in PD.

**Figure 5.**
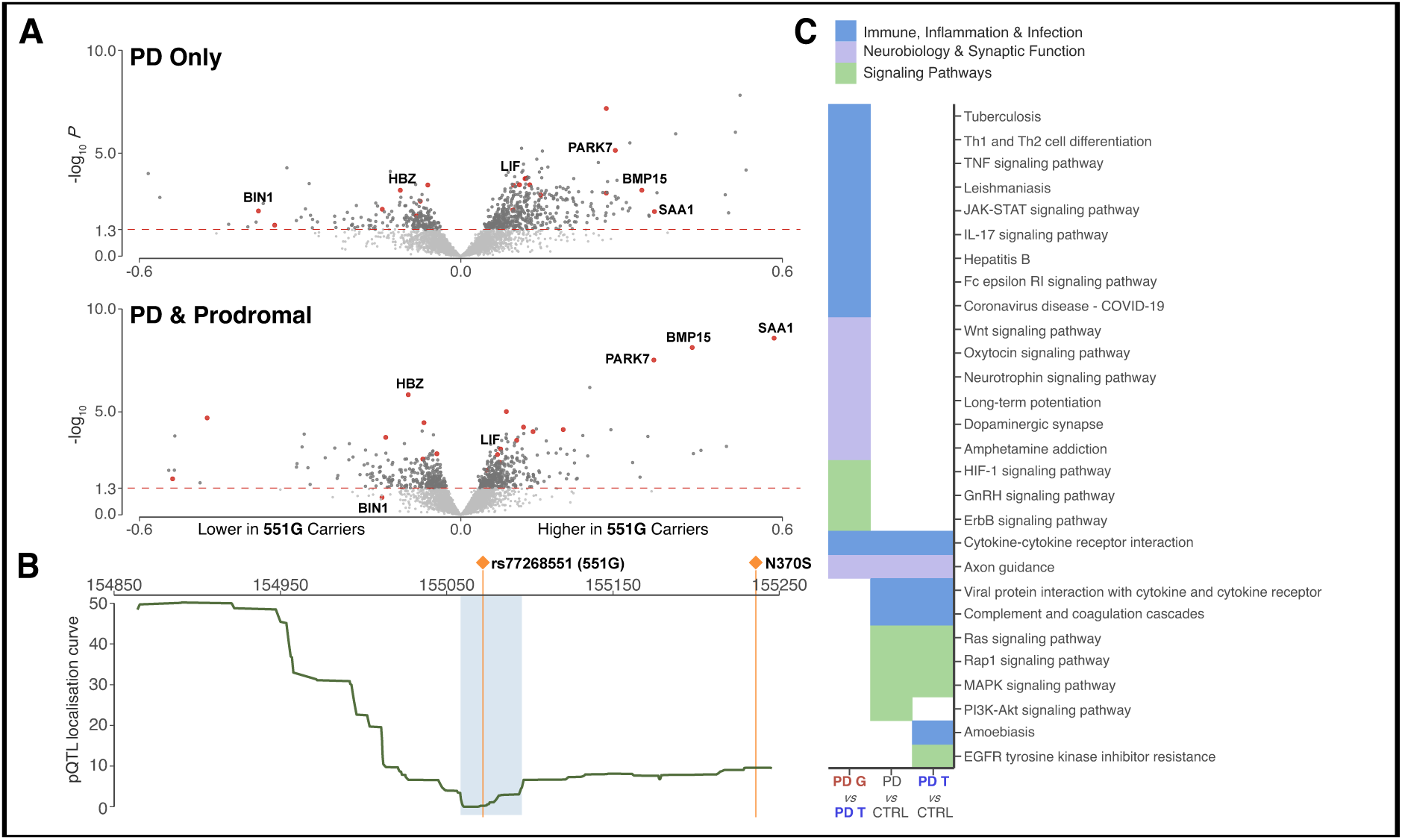
Allele-specific proteomic signatures and pathway mechanisms linked to 551G. **(A)** Volcano plots showing differential protein abundance between PD cases carrying the 551G risk allele and those carrying the alternative T allele (top = PD only; bottom = PD plus prodromal), combining data from both nEUR and AJ populations (black points). Proteins highlighted in red replicated independently across both populations, and labelled proteins additionally showed significant pQTL localisation within the narrow 551G interval. **(B)** meta-analysed pQTL localisation curve for the labelled proteins, demonstrating precise co-localisation of proteomic signals within the 551G locus. Individual pQTL localisation for each replicated protein are shown in the Supplementary Figure S10. **(C)** KEGG pathway enrichment across three contrasts: G versus T carriers, all PD cases versus controls, and T-only cases versus controls. Immune and infection-related pathways were enriched exclusively in the G versus T comparison, underscoring the specificity of the 551G-associated proteomic signature.

Notably, inclusion of prodromal cases increased the effect size of the SAA1 signal, supporting its potential as an early biomarker of disease activity. By contrast, BIN1 and HBZ were enriched in T-allele carriers, indicating divergence in downstream molecular profiles in non-carriers. BIN1, a protein implicated in Alzheimer’s disease^35^, showed reduced effect size upon inclusion of prodromal cases, consistent with a potential role in later-stage neurodegeneration rather than early inflammatory activation.

Finally, pathway enrichment analysis was conducted using KEGG (adjusted *P*<0.05) across three contrasts: i) G versus T carrier PD cases; ii) all PD cases versus controls, and iii) T-only carrier PD cases versus controls. Results revealed strong enrichment of immune and infection-related pathways exclusively in the G versus T comparison, including tuberculosis-related signalling (Figure 5C). The absence of comparable enrichment in the other contrasts underscores the specificity of the 551G-associated proteomic signature with *GBA1*-PD, while the distinct signalling profiles in T-only carriers, involving MAPK, RAP, and Ras pathways, point to the existence of other molecularly defined PD subtypes beyond the *GBA1*-PD haplotype.

## Discussion

What makes these findings compelling is not any single analysis, but the convergence of all of them. From large-scale genomic discovery across ∼900,000 IPDGC cases and controls, through transcriptomic, epigenomic, three-dimensional chromatin, and proteomic characterisation across two independent ancestries, every analytical approach independently maps to the same narrow kilobase interval distal to *GBA1*. These are not correlated readouts from a single technology, but orthogonal biological signals measured across independent omics platforms. This convergence defines a coherent mechanistic picture, implicating the 551G signal adjacent to *EFNA4* as the primary driver of the observed molecular effects, with *GBA1* itself emerging as a downstream regulated gene within this broader regulatory and haplotypic framework.

These findings refine the genetic architecture of the *GBA1* region and address a long-standing question in PD genetics by disentangling the relationship between *GBA1* and PD risk. Our results indicate that the apparent *GBA1* signal reflects the structure of an extended haplotype rather than a direct causal effect of protein-coding variation. Previous models have focused on loss- or gain-of-function effects of *GBA1*, implicitly treating it as the central driver of disease risk. In contrast, our findings suggest that the extended shared haplotype combines two distinct biological mechanisms, namely lysosomal enzyme deficiency and transcriptional dysregulation in sporadic PD, with both signals co-segregating on the same genetic background.

The evolutionary context of the extended haplotype provides additional perspective. Our signature of recent positive selection, replicated in population-based UK Biobank data, is consistent with a selective sweep spanning both N370S and the distal regulatory 551G. The latter represents a genetic hitchhiker, a passenger variant carried by the sweeping haplotype. A recent study^36^ reported that homozygous carriers of N370S exhibit increased resistance to tuberculosis (TB) potentially through sphingolipid accumulation within macrophage lysosomes that enhances bactericidal activity^37,38^. This effect was not observed in heterozygotes, favouring a model of homozygote protection against TBstudy^36^ as a strong selective force and challenging previous models of heterozygote advantage. Because N370S is a mild Gaucher variant^39^, and many (approximately two-thirds of) homozygotes remain asymptomatic with preserved fertility^37,40^, selection favouring tuberculosis resistance would have outweighed its modest fitness cost. Tuberculosis accounted for approximately half of deaths among Europeans of reproductive age during the late nineteenth century^41,42^, while AJ populations historically experienced repeated tuberculosis epidemics in densely populated European urban centres over many centuries. This prolonged exposure to tuberculosis likely imposed sustained positive selection favouring expansion of the extended shared haplotype that we observe today, carrying with it the 551G regulatory variant. This evolutionary linkage provides a compelling explanation for why association studies have focused on *GBA1* while overlooking the regulatory locus. It is precisely because selection shapes local LD structure that the LDU-based framework was uniquely positioned to capture this overlooked signal.

At the molecular level, the 551G locus is associated with a coordinated transcriptional programme involving eleven *cis*-regulated genes, including *GBA1* as a regulated target within a broader inflammatory network. Within this network, *GBA1* links transcriptional regulation to inflammatory biology through lysosomal stress, which is known to promote microglial activation and the release of pro-inflammatory cytokines, including IL-1β, IL-6, and TNFα^43^, providing a direct mechanistic connection between lysosomal dysfunction and neuroinflammation.

Beyond *GBA1*, the co-regulated target *cis-*genes support this network. Genes such as *IL6R, ADAR, SLC25A44, MEF2D, ARHGEF2, SEMA4A, GATAD2B*, and *ASH1L* implicate immune signalling, mitochondrial stress, RNA regulation, and chromatin remodelling. *IL6R* mediates innate immune activation and has been associated with neurodegeneration and cognitive decline in PD^44^, with pharmacological inhibition showing neuroprotective effects^45^. *ADAR* regulates RNA editing and modulates interferon responses^46^, while *SLC25A44* links mitochondrial metabolism to immune activation^47^. *ASH1L* suppresses proinflammatory cytokine production, and its dysregulation has been associated with chronic inflammation^48^. These transcriptional changes are also observed in a pericyte-specific context in our validation data (Figure 3C), reinforcing a link between immune activation and neuronal vulnerability.

A key question is how a single non-coding variant can exert such broad regulatory influence. We find that 551G resides within a neuronally active large enhancer exhibiting properties consistent with super-enhancer activity, including high transcription factor occupancy, enhancer RNA transcription, and characteristic histone modifications^49,50^. Supporting this interpretation, Hi-C and chromatin accessibility analyses indicate an allele-specific, hierarchical enhancer-promoter architecture in which the 551G enhancer connects to multiple distal regulatory elements before anchoring at the promoters of the *cis-*regulated genes. This nested interaction network, observed only in the carrier of the G risk allele and accompanied by increased chromatin accessibility, provides structural support for the proposed regulatory model. The evidence positions the enhancer as a regulatory hub^51^, propagating transcriptional activation in the region and linking genetic variation to coordinated transcriptional and protein output. Although 551G (rs77268551) represents the candidate causal variant at the best-fit localisation estimate, two flanking variants (rs11589479 and rs140309215) within the same support interval exhibit similar patterns of association. Functional fine-mapping will therefore be required to definitively resolve the causal variant within this narrow regulatory interval.

The downstream consequences of this regulatory architecture extend to the proteome. CSF analyses demonstrate that the same locus acts as a *trans*-pQTL, influencing protein abundance at distant genomic loci and bridging genetic risk with disease-relevant molecular phenotypes. This complements its role as a *cis*-eQTL for local gene expression, thus spanning transcriptional and proteomic levels. Among the identified *trans*-proteins, SAA1, PARK7, and BIN1 are particularly notable. SAA1, an acute-phase reactant, links systemic inflammation with central immune activation and is elevated in prodromal carriers, supporting its potential as an early biomarker^52^. PARK7 (DJ-1), a known monogenic PD gene^53,54^, is implicated here through a distinct regulatory mechanism, consistent with convergence between Mendelian and complex disease pathways and its role in modulating microglial activation and neuroinflammatory responses. By contrast, BIN1, enriched in 551 T-carriers, suggests divergence in downstream pathogenic pathways between genetically distinct PD subgroups that do not include *GBA1*-PD^35,55^.

Pathway analyses further reinforce these findings, revealing clear enrichment of immune and infection-related pathways specifically in 551G carriers, whereas non-carriers exhibit distinct signalling profiles involving MAPK, RAP, and Ras pathways. This contrast is important, as it demonstrates that the inflammatory programme observed in 551G carriers reflects a genetically defined disease mechanism rather than a generic feature of PD, while signalling profiles in non-carriers point to the presence of other distinct molecular subtypes of PD.

Convergence across independent genetic, evolutionary and multi-omic analyses supports a unifying framework in which 551G acts as the primary regulatory driver, inducing coordinated transcriptional and proteomic changes linked to neuroinflammatory pathways through chromatin-mediated remodelling. This primary role is supported directly by Hi-C data from the PD patient carrying 551G in the absence of N370S, showing that the distal regulatory element alone is sufficient to establish the allele-specific chromatin architecture independently of *GBA1* coding variation. We next asked whether this regulatory primacy was also reflected at the level of disease risk. To maximise sample size and haplotypic diversity, odds ratios were estimated from the fully phased haplotypes of the larger, pooled PPMI cohort (N = 864), comprising PD cases, prodromal individuals and controls, rather than the genetically homogeneous nEUR and AJ cohorts analysed throughout this study. The pooled cohort therefore captures a broader spectrum of European-derived haplotypes. Haplotypes carrying 551G alone and those carrying both 551G and N370S were compared with all remaining haplotypes carrying neither allele (Figure 6A).

**Figure 6.**
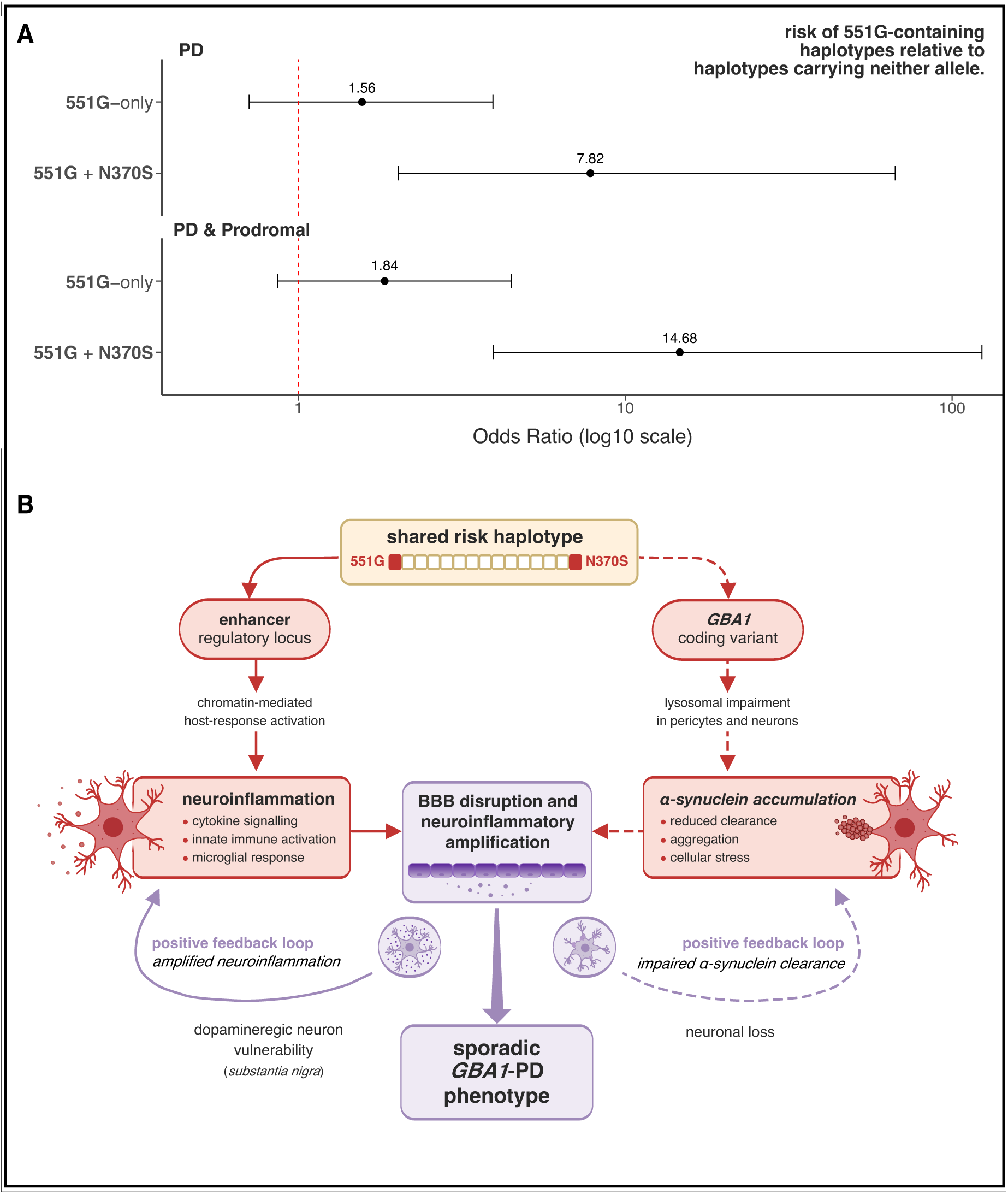
Dual-hit model for haplotype-driven PD. **(A)** Odds ratios for haplotypes carrying 551G alone or together with the *GBA1* N370S allele, compared against haplotypes carrying neither allele. N370S alone was almost never observed without 551G. **(B)** A shared 551G + N370S risk haplotype underlies the dual-hit model. The distal regulatory variant 551G acts as the primary driver of PD, inducing a coordinated neuroinflammatory programme through chromatin-mediated activation of host-response pathways. In parallel, N370S contributes to lysosomal dysfunction, impairing α-synuclein clearance. These complementary processes converge on blood-brain barrier (BBB) disruption and neuroinflammatory amplification. Positive feedback loops further reinforce both neuroinflammation and α-synuclein accumulation, promoting dopaminergic neuron vulnerability and neuronal loss. The interplay of these genetic and molecular mechanisms drive the clinically distinct *GBA1*-PD phenotype.

The N370S-only comparison was not estimable because N370S was almost never observed without 551G, indicating that the previous IPDGC GWAS analyses were capturing the shared 551G+N370S haplotype rather than the effect of N370S in isolation. Haplotypes carrying 551G alone conferred a PD odds ratio of 1.56, increasing to 1.84 when prodromal cases were included, providing further genetic evidence that 551G contributes to disease susceptibility independently of *GBA1* coding variation. In contrast, co-inheritance of N370S on the same haplotype amplified the odds ratio to 7.82 for PD and to 14.68 when prodromal cases were included, consistent with a synergistic dual-hit architecture in which N370S modifies and amplifies the risk established by 551G but cannot operate without it.

All our complementary analyses distinguish the primary regulatory contribution of 551G from the risk-amplifying effect of co-inherited *GBA1* coding variation. On this extended haplotypic background, upregulation of *GBA1* observed in blood does not necessarily imply preserved or increased enzymatic activity. Instead, it likely reflects a compensatory transcriptional response to lysosomal stress, consistent with known feedback mechanisms in lysosomal biology. In this context, the N370S variant is expected to impair lysosomal function^36^ or GCase folding and trafficking, resulting in reduced α-synuclein clearance. Distal regulatory variation and *GBA1*-related functional effects therefore act together to influence disease-relevant inflammatory pathways. These findings support a dual-hit model, illustrated in Figure 6B, in which a permissive regulatory environment and *GBA1*-related functional impairment act synergistically, such that α-synuclein accumulation alone is not sufficient to drive pathology but becomes pathogenic in the presence of sustained inflammatory stress.

This model is further supported by cell-type-specific evidence. Pericytes and microglia play key roles in α-synuclein clearance via lysosomal pathways^56^. Previous work^56^ has shown that α-synuclein accumulation alone does not induce inflammation, but becomes cytotoxic only in the presence of an additional cellular insult, such as inflammatory activation. The co-occurrence of genetic variation affecting lysosomal function and inflammatory signalling on the same haplotypic background provides a plausible mechanistic basis for disease initiation in this subgroup of PD patients, potentially involving disruption of blood-brain barrier integrity and amplification of neuroinflammation feedback loops (Figure 6B).

To conclude, our findings support a multi-component model of genetic risk, rather than a single-variant mechanism, in which regulatory and coding variants co-segregate on a shared haplotype and act in concert. Our data indicate that the distal enhancer is sufficient to drive PD-associated molecular changes, supported by convergent evidence of neuroinflammatory pathways, whereas co-inheritance with *GBA1* coding variation modulates disease expression and gives rise to the clinically distinct *GBA1*-PD phenotype. While the shared risk haplotype carrying both variants is the most common configuration in the populations studied, further work in controlled cellular systems will be required to define how these components interact and to establish their clinical relevance across diverse genetic backgrounds.

Multiple clinical trials are currently underway to target *GBA1* coding variation, yet our findings situate *GBA1* within a broader regulatory and neuroinflammatory framework that may require a different therapeutic perspective. By integrating population genetics with chromatin architecture and multi-omics mapping, this study demonstrates how complex haplotypic architecture can obscure causal mechanisms using conventional mapping approaches, and how modelling population-specific LD structure directly within the localisation framework can resolve biological complexity that has resisted standard analytical strategies. The depth and harmonised design of the PPMI dataset further illustrate the value of disease-specific, deeply phenotyped resources for dissecting complex inheritance, and provide a template for applying integrative multi-omics frameworks across complex diseases. More broadly, our findings support the emerging view that in neurodegeneration, genetic risk operates through the convergence of regulatory, inflammatory, and cell biological mechanisms.

## ACKNOWLEDGMENTS

This study was funded by the Medical Research Council (MRC UK; MR/X011070/1). The authors gratefully acknowledge the landmark PPMI study (www.ppmi-info.org) for providing access to the data and all its study participants, the FOUNDIN-PD project (www.foundinpd.org) for providing access to the multi-layered molecular data from PPMI iPSC-derived dopaminergic neurons, and the UK Biobank for population-based data (www.ukbiobank.ac.uk). The authors also acknowledge the UCL Computer Science High Performance Computing Cluster (https://hpc.cs.ucl.ac.uk/) for providing the computational resources. This work is dedicated to the memory of Prof Newton E. Morton (1931–2018), whose pioneering contributions to human genetics inspired the earlier methodological work that laid the foundation for this study. <u>Official PPMI</u> <u>acknowledgement:</u> Data used in the preparation of this article was obtained on [2023-11-10] from the Parkinson’s Progression Markers Initiative (PPMI) database (www.ppmi-info.org/access-dataspecimens/download-data), RRID:SCR_006431. For up-to-date information on the study, visit www.ppmi-info.org. PPMI – a public-private partnership – is funded by the Michael J. Fox Foundation for Parkinson’s Research and funding partners, including 4D Pharma, Abbvie, AcureX, Allergan, Amathus Therapeutics, Aligning Science Across Parkinson’s, AskBio, Avid Radiopharmaceuticals, BIAL, BioArctic, Biogen, Biohaven, BioLegend, BlueRock Therapeutics, Bristol-Myers Squibb, Calico Labs, Capsida Biotherapeutics, Celgene, Cerevel Therapeutics, Coave Therapeutics, DaCapo Brainscience, Denali, Edmond J. Safra Foundation, Eli Lilly, Gain Therapeutics, GE HealthCare, Genentech, GSK, Golub Capital, Handl Therapeutics, Insitro, Jazz Pharmaceuticals, Johnson & Johnson Innovative Medicine, Lundbeck, Merck, Meso Scale Discovery, Mission Therapeutics, Neurocrine Biosciences, Neuron23, Neuropore, Pfizer, Piramal, Prevail Therapeutics, Roche, Sanofi, Servier, Sun Pharma Advanced Research Company, Takeda, Teva, UCB, Vanqua Bio, Verily, Voyager v. 25MAR2024 Therapeutics, the Weston Family Foundation and Yumanity Therapeutics.

